# Development and validation of a six-RNA binding proteins prognostic signature and candidate drugs for prostate cancer

**DOI:** 10.1101/2020.06.28.175984

**Authors:** Lei Gao, Jialin Meng, Yong Zhang, Junfei Gu, Zhenwei Han, Shenglin Gao, Xiaolu Wang

## Abstract

The dysregulation of RNA binding proteins (RBPs) play critical roles in the progression of several cancers. However, the overall functions of RBPs in prostate cancer (PCa) remain poorly understood. Therefore, we first identified 144 differentially expressed RBPs in tumors compared to normal tissues based on the TCGA dataset. Next, six RBP genes (MSI1, MBNL2, LENG9, REXO2, RNASE1, PABPC1L) were screened out as prognosis hub genes by univariate, LASSO and multivariate Cox regression and used to establish the prognostic signature. Further analysis indicated that high risk group was significantly associated with poor RFS, which was validated in the MSKCC cohort. Besides, patients in high risk group was closely associated with dysregulation of DNA damage repair pathway, copy number alteration, tumor burden mutation and low-respond to cisplatin (P < 0.001), bicalutamide (P < 0.001). Finally, three drugs (ribavirin, carmustine, carbenoxolone) were predicted using Connectivity Map. In summary, we identified a six-RBP gene signature and three candidate drugs against PCa, which may promote the individualized treatment and further improve the life quality of PCa patients.

## 1. Introduction

Prostate cancer (PCa) is the most common cancer among men, and one of the leading cause of cancer-specific deaths. In the United States, the projected number of new cases in 2020 is about 191,930, while the estimated number of deaths is high to 33,330, only less than lung cancer in men[1]. In China, the incidence of PCa has risen rapidly from 10% to 20% over the past two decades[2]. Though the overall survival of PCa is not poor compared to other malignancies[3-5], the recurrence rate of it is very high, and most patients will develop into an advanced stage of so called castration-resistant PCa (CRPC) [6, 7]. Moreover, patients with metastatic status only have a short median survival of 30 months[8]. Therefore, further exploring the molecular mechanism of PCa, and develop effective methods for screening and diagnosis are pivotal to improve life quality for patients.

RNA binding proteins (RBPs) are a group of proteins which could function with the RNA-binding domain to distinguish and bind to target RNAs, including coding RNA and non-coding RNAs (rRNAs, ncRNAs, snRNAs, miRNAs, tRNAs, and snoRNAs)[9]. Up to now, more than 1,500 RBP genes have been founded through genome-wide screening in human genome[10]. RBPs could regulate gene transcription, but mainly act on RNA processing, such as mRNA splicing, localization, polyadenylation, translocation, stability, translation[11, 12], which is more rapid than transcription factors mediated regulation[13, 14]. Due to the critical roles of post-transcriptional regulation in the development of biology, it is thus reasonable that functional disruption of RBPs are involved into the progression of multiple human diseases, such as U1-70K in Alzheimer’s disease[15], muscleblind in neurodegeneration[16, 17], FMRP in Fragile X syndrome [18], quaking (QKI), human antigen R (HuR), and serine/arginine-rich splicing factor 1 (SRSF1) in cardiovascular disease[19]. Even though RBPs are known to be involved in the biology and progression of numerous diseases, the roles of RBPs in carcinoma development are still rare.

In the past decades, it has been reported that many RBPs were aberrantly changed in tumors, which affected the procession of mRNA into protein level, and then involved in carcinogenesis[20, 21]. For example, AGO2 promotes tumor progression via involving into the oncogenic miR-19b biogenesis[22]; ADAR1 facilitates the tumorigenesis of PCa by regulating the expression and stability of the prostate cancer antigen 3 (PCA3)[23, 24]; dysregulated expression of IGF2BP3 accelerates gastric carcinogenesis due to the silence of miR-34a[25]; QKI-5 regulates cell proliferation in lung cancer by changing cancer-associated alternative splicing[26].In general, these results demonstrate that the RBPs are closely associated with the progression of human tumors. Therefore, we collected PCa expression profiles and clinical information from the cancer genome atlas (TCGA) database. Then, differentially expressed RBPs between tumor and normal tissues were filtered, and the systematic analysis to seek out the potential mechanisms and clinical information of RBPs were also performed. Several differentially expressed RBPs involved with PCa were identified, which may provide novel thought for the progression of PCa.

## 2. Materials and methods

### 2.1. Data processing

The flowchart of our study was displayed in Figure 1, the RNA-sequencing dataset (FPKM value) of gene expression of 52 normal prostate tissue samples and 499 PCa samples with related clinical data were downloaded from the Genomic Data Commons (GDC, https://portal.gdc.cancer.gov/). Then FPKM values were transformed into transcripts per kilobase million (TPM) values. The mRNA expression profiles of the 1452 RBPs were obtained from archive of the PCa project. To identify the differently expressed RBPs in view of |log2 fold change (FC)|>0.5 and false discovery rate (FDR) < 0.05. The MSKCC cohort was also included as validation cohorts to evaluate the prognostic ability of RBPs.

**Figure 1.**
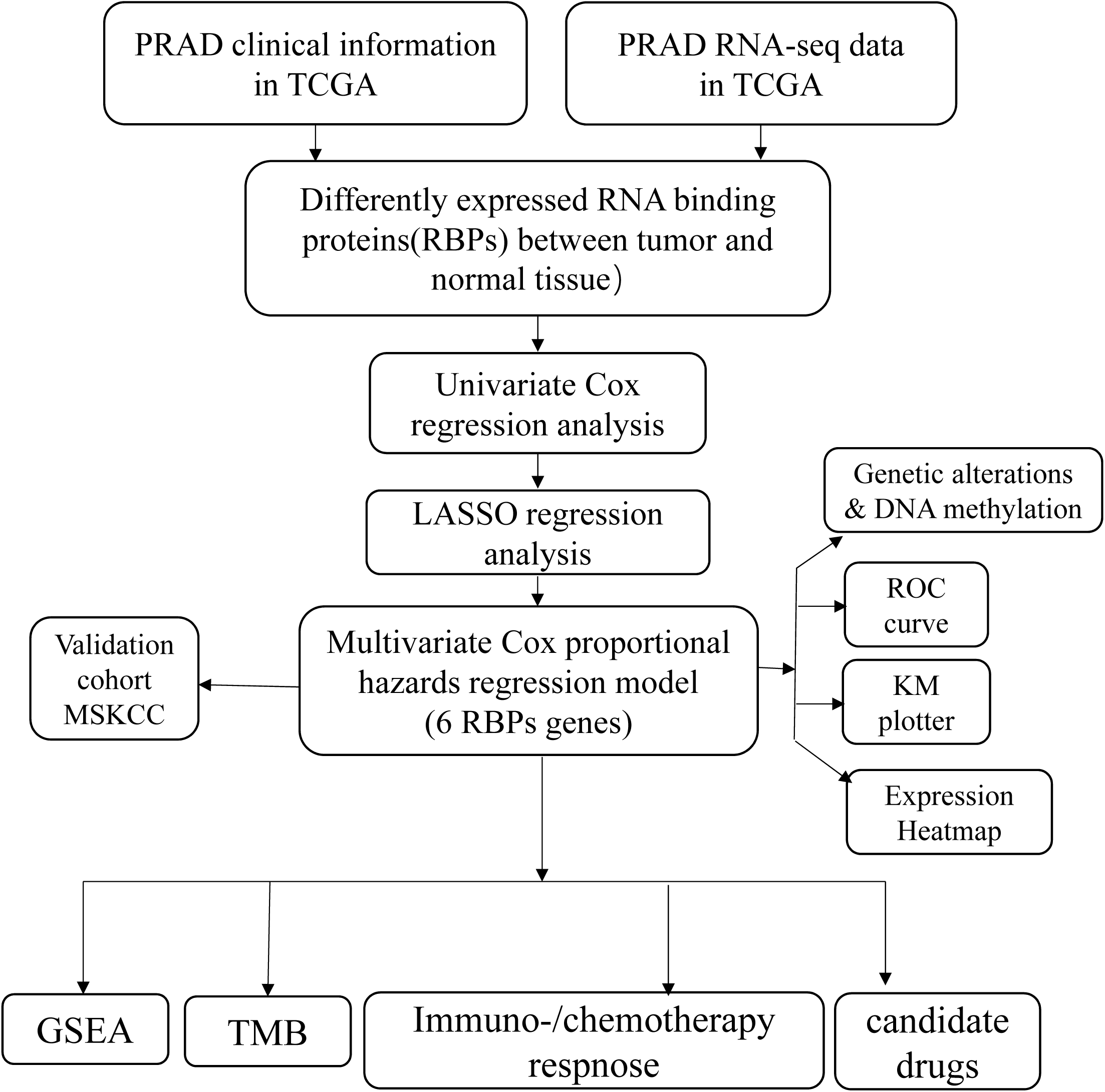
Flowchart for analyzing RBPs in prostate cancer.

### 2.2. Prognosis-Related hub RBPs detection and validation

The differentially expressed RBPs were subjected to a univariate Cox regression analysis using the survival package in R (https://github.com/therneau/survival). Next, the least absolute shrinkage and selection operator (LASSO)[27], was performed to further filter the prognostic RBPs candidates. In addition, multiple stepwise Cox regression method was used to select the hub RBPs to construct the prognostic model. Risk score of selected hub genes was calculated based on the mRNA expression weighted by the estimated regression coefficient in the multiple cox regression analysis.

### 2.3. Gene expression, genetic alteration, DNA methylation analysis of hub RBPs

The genetic alterations and DNA methylation data of hub RBP genes in PCa patients were collected from the cBioPortal platform (http://www.cbioportal.org/). OncoPrinter were performed to display the frequency of genetic alteration, correlation between copy number alteration (CNV), methylation and mRNA expression were calculated and visualized by ggplot2 R package (https://github.com/tidyverse/ggplot2).

### 2.4. Identification of risk-associated Different Expression Genes (DEGs) and enrichment analysis

The PCa patients were divided into high and low risk group based on the risk score, and DEGs were identified by using R package ‘limma’[28]. Then Gene Ontology (GO) analysis of these DEGs were performed by utilizing clusterProfiler R package[29]. GSEA[30] was applied to explore the potential activated pathways in high risk group of RBPs. Annotated gene sets h.all.v7.1.symbols.gmt, c2.cp.kegg. v7.1.symbols.gmt were selected as the reference gene sets.

### 2.5. DNA damage repair (DDR) pathway, tumor mutation burden (TMB) and tumor stemness

The mRNA expression and genetic alterations of 21 hub gene from DDR pathway[31] between different risk group were detected based on the TCGA cohort. TMB, which is defined as the total number of mutations per megabyte of tumor tissue, was calculated by dividing the total number of mutations by the size of the coding region of the target. Tumor stemness, could be represented using mRNAsi, then we got the mRNAsi data of TCGA PCa patients from Robertson’s report[32].

### 2.6. Immunotherapy response, chemotherapy response prediction and candidate small molecules

The Tumour Immune Dysfunction and Exclusion (TIDE) (http://tide.dfci.harvard.edu) was performed to assess the individual likelihood of responding to immunotherapy. The chemotherapy response for cisplatin, docetaxel and bicalutamide of each PCa patient in TCGA cohort was calculated based on the Genomics of Drug Sensitivity in Cancer (GDSC, https://www.cancerrxgene.org) by using ‘pRRophetic’ R package (https://github.com/paulgeeleher/pRRophetic). To identify potential drugs for RBPs risk group, Connectivity map (CMap)[33] was performed to predict small molecule drugs for RBPs high risk group compared to RBPs low risk group.

### 2.7. Statistics

All data were performed in R platform (v.3.6.3, https://cran.r[project.org/). The comparison of mRNA levels of RBPs between PCa and normal tissues, CNV, TMB, chemotherapy response, immunotherapy response, tumor stemness of different risk groups were conducted using Wilcoxon rank sum tests. The hub RBPs were screened out using Cox regression, LASSO and multiple stepwise Cox regression. The Kaplan-Meier (K-M) curve was generated using the R package ‘survival’ and assessed by log-rank test. The risk score of each PCa patient was calculated using the mRNA expression of hub RBPs, weighted by the corresponding coefficients derived from multiple Cox regression model.

## 3. Results

### 3.1. Explore the prognostic RBPs in PCa

A total of 499 PCa samples and 52 normal prostate samples were analyzed. The limma software package were used to preprocess these data and detect the differentially expressed RBPs. In total, 1542 RBPs[10] were included in this study, of which 144 met our threshold (adj P < 0.05, |log2FC|>0.5), 51 downregulated and 93 upregulated RBPs (Figure 2A, 2B).

**Figure 2.**
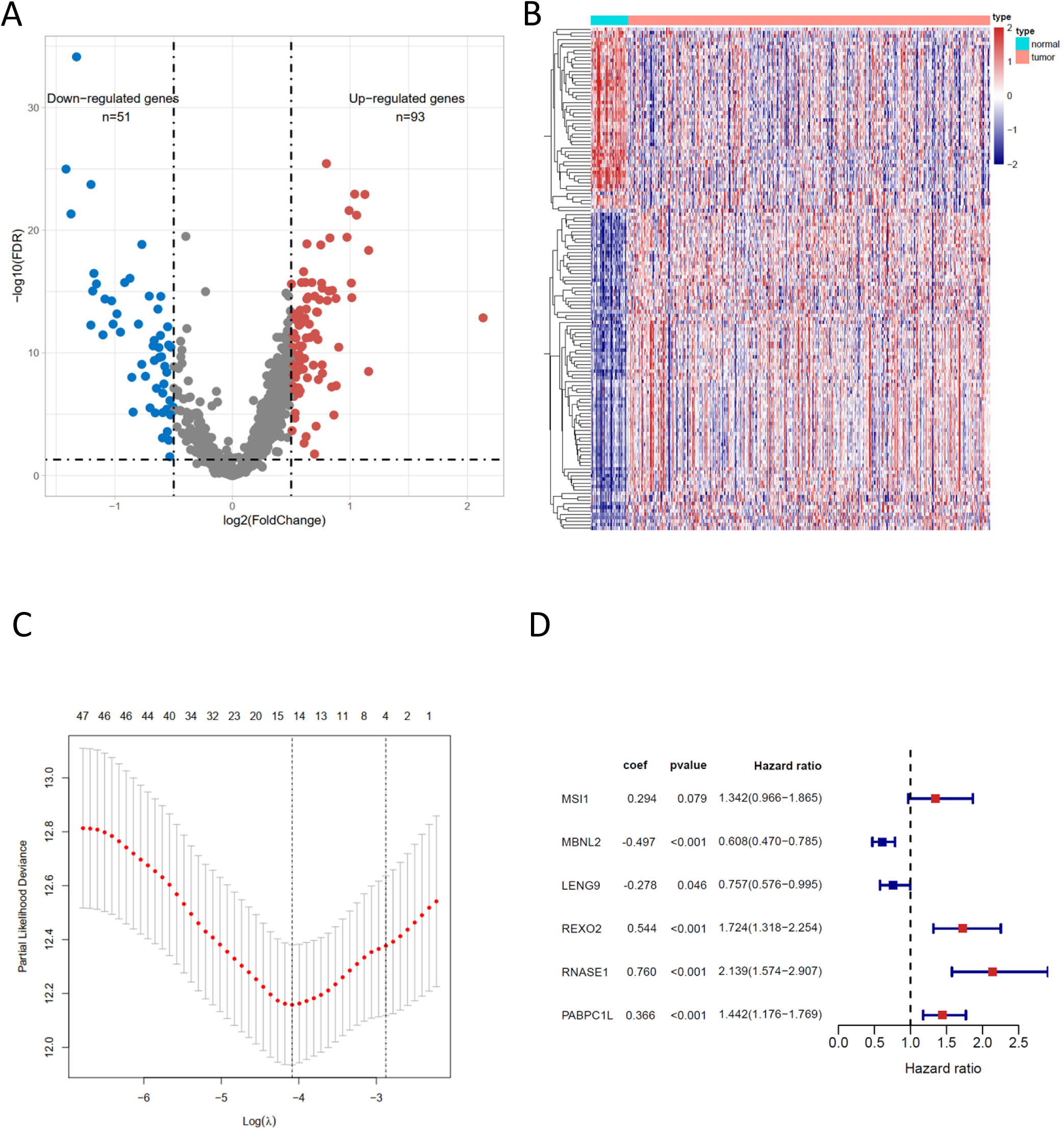
The differentially expressed RBPs, lasso regression results and risk prognostic model of RBPs. (A) Volcano plot represents the differentially expressed RBPs, satisfying the criteria of adjusted pvalue < 0.05, |log2FoldChange| > 0.5. (B) Heatmap plot represents the expressions of 144 differentially RBPs between normal and tumor samples in prostate cancer. (C) The Cross-Validation fit curve calculated by lasso regression method. (D) The coefficients of six genes estimated by multivariate Cox regression.

The 144 differently expressed RBPs were included to investigate the prognostic significance by using univariate Cox regression method and obtained 49 prognostic-associated candidate RBPs (**Table 1**). Then LASSO regression was performed to identify the 49 candidate genes that were closely associated with the prognosis of PCa patients, including 14 genes, PPARGC1A, EZH2, MSI1, MBNL2, SNRNP70, LENG9, PABPN1, REXO2, XPO6, RNASE1, PABPC1L, LUC7L, SMAD9, ZFP36 (**Figure 2C**). To further select the RBPs with the greatest prognosis value, multiple stepwise Cox regression were performed to investigate their impact, six hub RBPs were selected to construct the risk model in PRAD patients (**Figure 2D**). The expressions profiles of these six genes showed that MSI1, LENG9, REXO2, PABPC1L were overexpression in prostate cancer samples when compared with normal samples, and MBNL2, RNASE1 were lower in tumor tissue (**Figure 3A**). Moreover, the K-M plot also demonstrated that overexpression of MSI1, REXO2, RNASE1, PABPC1L, and low expression of MBNL2, LENG9 were associated with the poor RFS of PCa patients (All, *P* <0.001) (**Figure 3B**). Finally, the genetic alteration status and methylation of six RBPs was explored based on cBioPortal database. The results showed that the MSI1, MBNL2, LENG9, REXO2, RNASE1, PABPC1L had 1.8%, 2.2%, 0.6%, 4%, 0.4%, 0.6% genetic alterations (**Figure 4A**), and most of them belonged to copy number change. The correlation between copy number mutation with mRNA expression of MSI1 was 0.376 (Figure 4B), the other genes were less than 0.3. What’s more, the correlation between mRNA expression and DNA methylation of MBNL2, REXO2, RNASE1 were −0.470, −0.640, −0.486 (**Figure 4C**), which suggested that MSI1 may be copy number-drive gene, while MBNL2, REXO2, RNASE1 may be methylation-drive genes.

**Table 1.** Unicox results of differential RBPs.

| gene | HR | HR.95L | HR.95H | pvalue |
| --- | --- | --- | --- | --- |
| PPARGC1A | 0.521 | 0.356 | 0.764 | 0.001 |
| POLR2H | 2.434 | 1.567 | 3.782 | 0 |
| RBMS3 | 0.711 | 0.536 | 0.945 | 0.019 |
| EZH2 | 2.081 | 1.533 | 2.826 | 0 |
| MSI1 | 1.852 | 1.313 | 2.612 | 0 |
| GNL3 | 1.593 | 1.072 | 2.366 | 0.021 |
| NOP16 | 1.722 | 1.177 | 2.52 | 0.005 |
| MBNL2 | 0.727 | 0.583 | 0.906 | 0.005 |
| PPAN | 1.416 | 1.003 | 2 | 0.048 |
| MRPL41 | 1.391 | 1.045 | 1.851 | 0.024 |
| QTRT1 | 1.761 | 1.249 | 2.485 | 0.001 |
| GNL2 | 1.814 | 1.048 | 3.142 | 0.033 |
| MEX3D | 1.635 | 1.13 | 2.365 | 0.009 |
| SNRNP70 | 1.944 | 1.427 | 2.648 | 0 |
| RRP12 | 1.596 | 1.011 | 2.519 | 0.045 |
| LENG9 | 0.754 | 0.584 | 0.972 | 0.029 |
| TARBP1 | 1.739 | 1.289 | 2.346 | 0 |
| MTG1 | 1.803 | 1.295 | 2.51 | 0 |
| RUVBL1 | 1.787 | 1.233 | 2.59 | 0.002 |
| PUS1 | 2.061 | 1.381 | 3.075 | 0 |
| RRS1 | 1.52 | 1.085 | 2.129 | 0.015 |
| RPL28 | 1.406 | 1.002 | 1.975 | 0.049 |
| TRMT1 | 2.063 | 1.372 | 3.102 | 0.001 |
| PABPN1 | 3.232 | 2.008 | 5.202 | 0 |
| TRMU | 2.332 | 1.559 | 3.488 | 0 |
| METTL1 | 1.839 | 1.102 | 3.067 | 0.02 |
| SNRPF | 2.451 | 1.636 | 3.674 | 0 |
| RRP9 | 2.245 | 1.404 | 3.59 | 0.001 |
| RPL38 | 1.504 | 1.071 | 2.114 | 0.019 |
| REPIN1 | 1.849 | 1.213 | 2.819 | 0.004 |
| RPL18A | 1.407 | 1.023 | 1.935 | 0.036 |
| TRMT2A | 1.855 | 1.177 | 2.922 | 0.008 |
| LSM7 | 1.634 | 1.183 | 2.256 | 0.003 |
| RPL37A | 1.525 | 1.072 | 2.169 | 0.019 |
| RPS28 | 1.521 | 1.114 | 2.076 | 0.008 |
| RPS29 | 1.557 | 1.1 | 2.203 | 0.013 |
| RPUSD1 | 1.764 | 1.194 | 2.606 | 0.004 |
| REXO2 | 1.592 | 1.245 | 2.035 | 0 |
| RPL31 | 1.607 | 1.063 | 2.429 | 0.024 |
| XPO6 | 1.52 | 1.137 | 2.032 | 0.005 |
| MEX3A | 1.181 | 1.009 | 1.383 | 0.039 |
| RNASE1 | 1.39 | 1.057 | 1.829 | 0.019 |
| TIPARP | 0.762 | 0.606 | 0.959 | 0.02 |
| PABPC1L | 1.608 | 1.323 | 1.955 | 0 |
| LUC7L | 2.322 | 1.67 | 3.228 | 0 |
| RPS21 | 1.431 | 1.055 | 1.942 | 0.021 |
| SMAD9 | 0.788 | 0.625 | 0.993 | 0.044 |
| ANG | 0.826 | 0.693 | 0.984 | 0.033 |
| ZFP36 | 0.83 | 0.716 | 0.962 | 0.013 |

**Figure 3.**
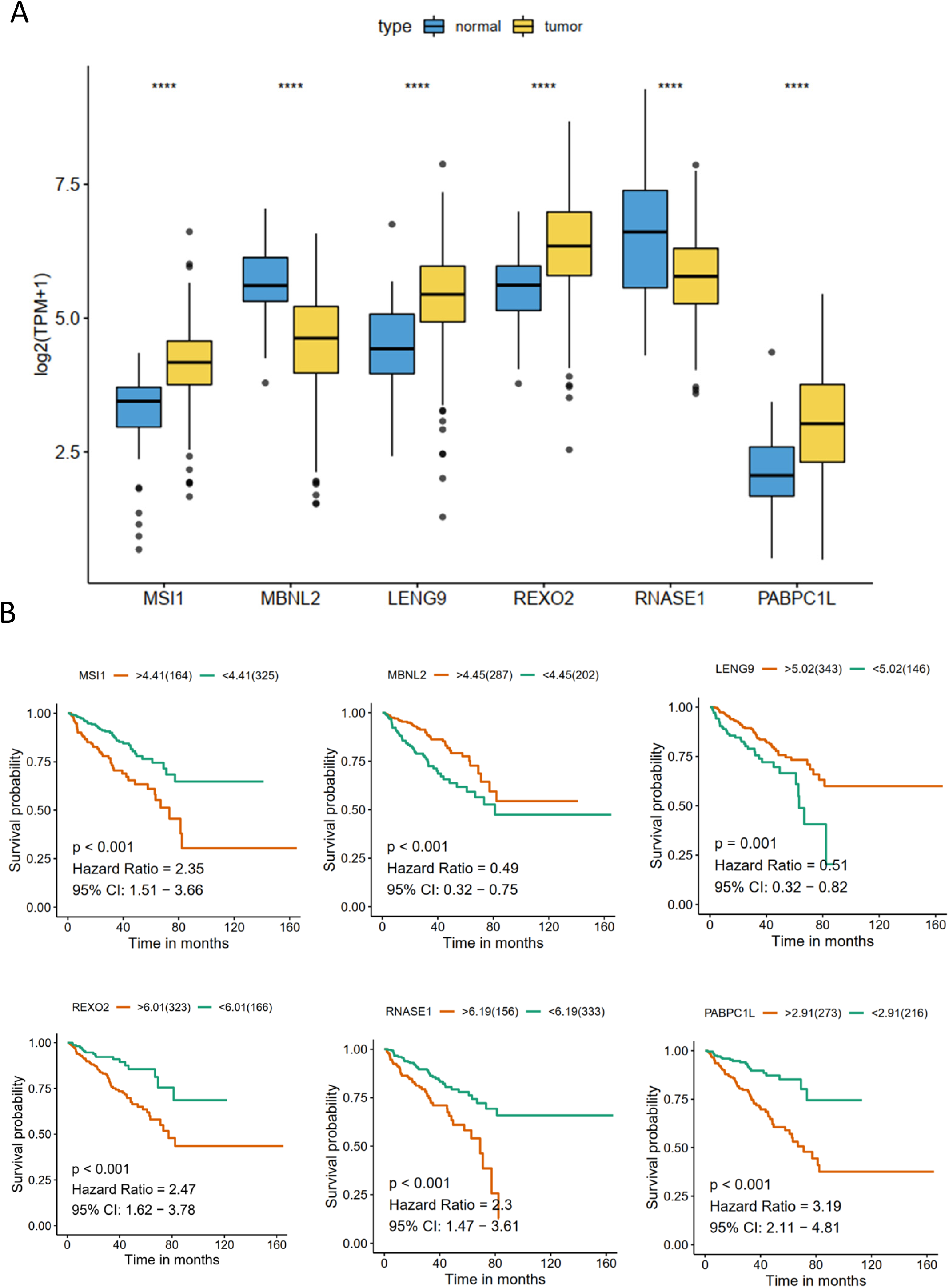
The mRNA expression profiles, survival curves of six hub RBPs. (A) The mRNA expression profiles of MSI1, MBNL2, LENG9, REXO2, RNASE1, PABPC1L. (B) The survival curves of MSI1, MBNL2, LENG9, REXO2, RNASE1, PABPC1L.

**Figure 4.**
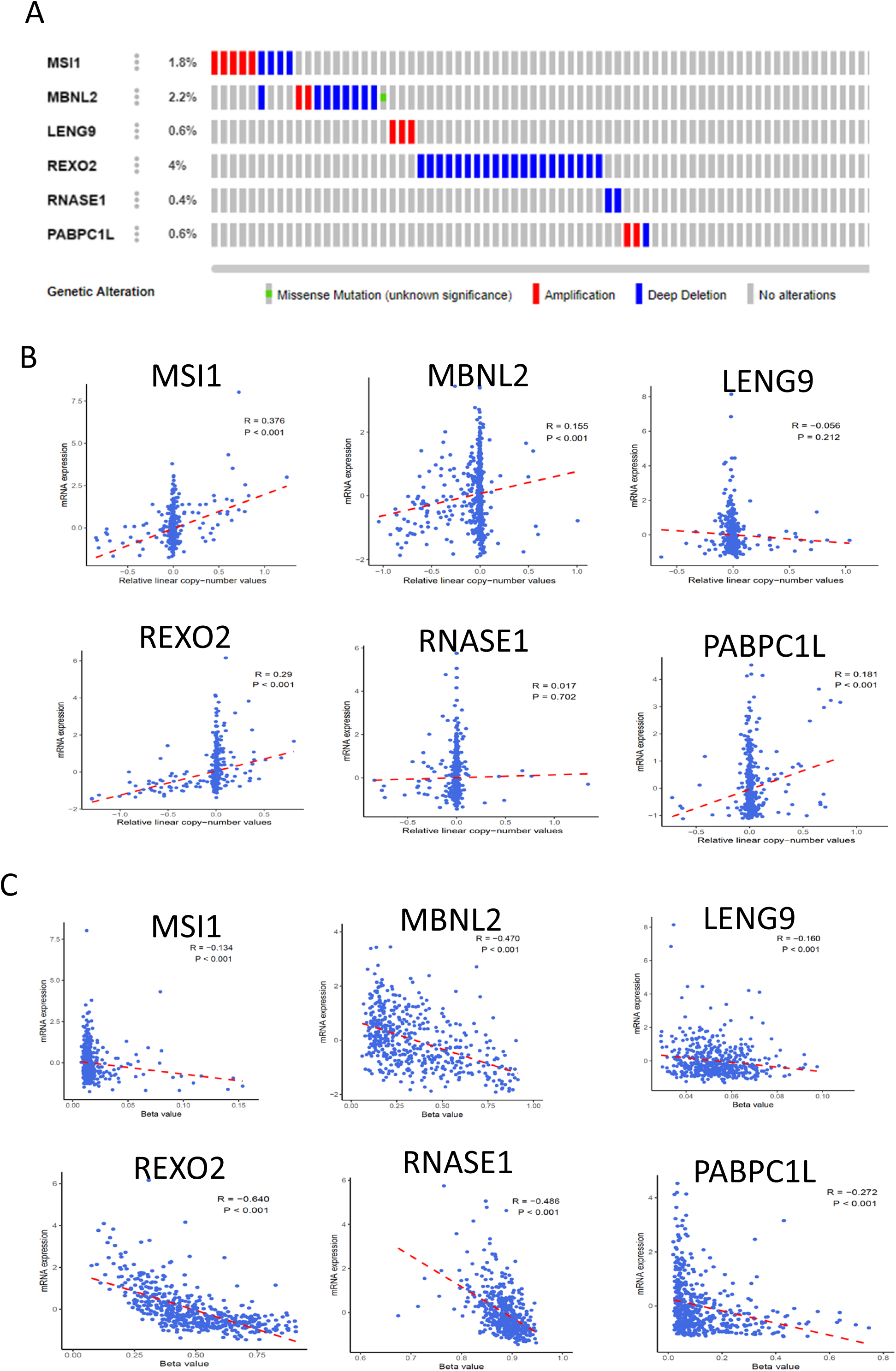
The genetic alterations and methylation of the six hub RBPs. (A) The genetic alterations of the six genes. (B) The correlation between mRNA expression and copy number values defined by GISTIC2 method. (C) The correlation between mRNA expression and DNA methylation of the six genes.

### 3.2. Construction and validation of the RBPs prognostic model

The six hub RBPs choosed from the multiple Cox regression method were used to establish the predictive model. The risk score of each patient was calculated based on the coefficients: Risk score = (0.294*ExpMSI1) + (−0.497*ExpMBNL2) + (−0.278*ExpLENG9) + (0.544*ExpREXO2) + (0.760*ExpRNASE1) + (0.366*ExpPABPC1L). Then PCa patients were divided into low-risk group (n =245) and high-risk group (n=244) based on the median risk score as the cut-off point. The K-M results showed that the patients in the high-risk group had worse RPS compared to those in the low-risk group (*P* < 0.001, **Figure 5A**). Moreover, the time-dependent ROC demonstrated that the area under the ROC curve (AUC) of this RBPs risk score model in 1-year, 3-year, 5-year could be 0.799, 0.736, 0.714 (Figure 5B), which indicated that it has moderate diagnostic performance. The RFS status of patients, heatmap of expression profiles of these six genes in the low-risk group and high-risk group were displayed **Figure 5C** and **Figure 5D**. Moreover, the univariate and multivariate Cox regression analysis results showed that risk score was an independent factor for RFS (**Table 2**).To verify whether the six RBPs model with similar predictive value in other PCa cohort, the same formula was used to generate the risk score of each patient in the MSKCC dataset. Consistent with the results in the TCGA cohort, patients with high-risk score also have a not good RFS than those with low-risk score (**Figure 6A–6D**).

**Table 2.** Unicox and multicox regression analysis of riskscore and clinical informations.

| Variables | unicox |  |  | multicox |  |  |
| --- | --- | --- | --- | --- | --- | --- |
|  | HR | 95%CI of HR | *P* | HR | 95%CI of HR | *P* |
| age | 1.03 | 0.99-1.06 | 0.055 | 1.01 | 0.98-1.05 | 0.419 |
| gleason | 2.23 | 1.80-2.77 | <0.001 | 1.89 | 1.48-2.41 | <0.001 |
| T | 2.54 | 2.54-1.69 | <0.001 | 1.09 | 1.09-0.67 | 0.727 |
| riskScore | 1.25 | 1.25-1.19 | <0.001 | 1.19 | 1.12-1.26 | <0.001 |

**Table 3.** Results of CMap analysis.

| cmap name | mean | n | enrichment | p | specificity | percent<br>non-null |
| --- | --- | --- | --- | --- | --- | --- |
| **ribavirin** | -0.386 | 4.000 | -0.860 | 0.001 | 0.000 | 50.000 |
| **carmustine** | -0.632 | 3.000 | -0.805 | 0.015 | 0.034 | 66.000 |
| **carbenoxolone** | -0.037 | 4.000 | -0.697 | 0.018 | 0.013 | 50.000 |
| famotidine | -0.462 | 5.000 | -0.564 | 0.048 | 0.023 | 60.000 |
| tridihexethyl | -0.383 | 4.000 | -0.490 | 0.202 | 0.328 | 50.000 |
| butoconazole | -0.390 | 4.000 | -0.473 | 0.238 | 0.252 | 50.000 |
| metergoline | -0.165 | 4.000 | -0.459 | 0.268 | 0.200 | 50.000 |
| NU-1025 | -0.331 | 2.000 | -0.621 | 0.290 | 0.247 | 50.000 |
| mercaptopurine | -0.319 | 2.000 | -0.547 | 0.411 | 0.432 | 50.000 |
| STOCK1N-35874 | 0.434 | 2.000 | 0.501 | 0.497 | 0.571 | 50.000 |
| 5279552 | 0.432 | 2.000 | 0.492 | 0.529 | 0.528 | 50.000 |
| AH-6809 | -0.433 | 2.000 | -0.491 | 0.533 | 0.592 | 50.000 |
| STOCK1N-35696 | 0.424 | 2.000 | 0.490 | 0.537 | 0.729 | 50.000 |
| 5255229 | -0.426 | 2.000 | -0.489 | 0.542 | 0.418 | 50.000 |
| 5230742 | -0.416 | 2.000 | -0.486 | 0.552 | 0.656 | 50.000 |
| imatinib | 0.270 | 2.000 | 0.456 | 0.660 | 0.796 | 50.000 |
| MK-886 | 0.254 | 2.000 | 0.455 | 0.663 | 0.708 | 50.000 |
| TTNPB | 0.253 | 2.000 | 0.455 | 0.664 | 0.776 | 50.000 |
| exisulind | 0.227 | 2.000 | 0.454 | 0.665 | 0.652 | 50.000 |

**Figure 5.**
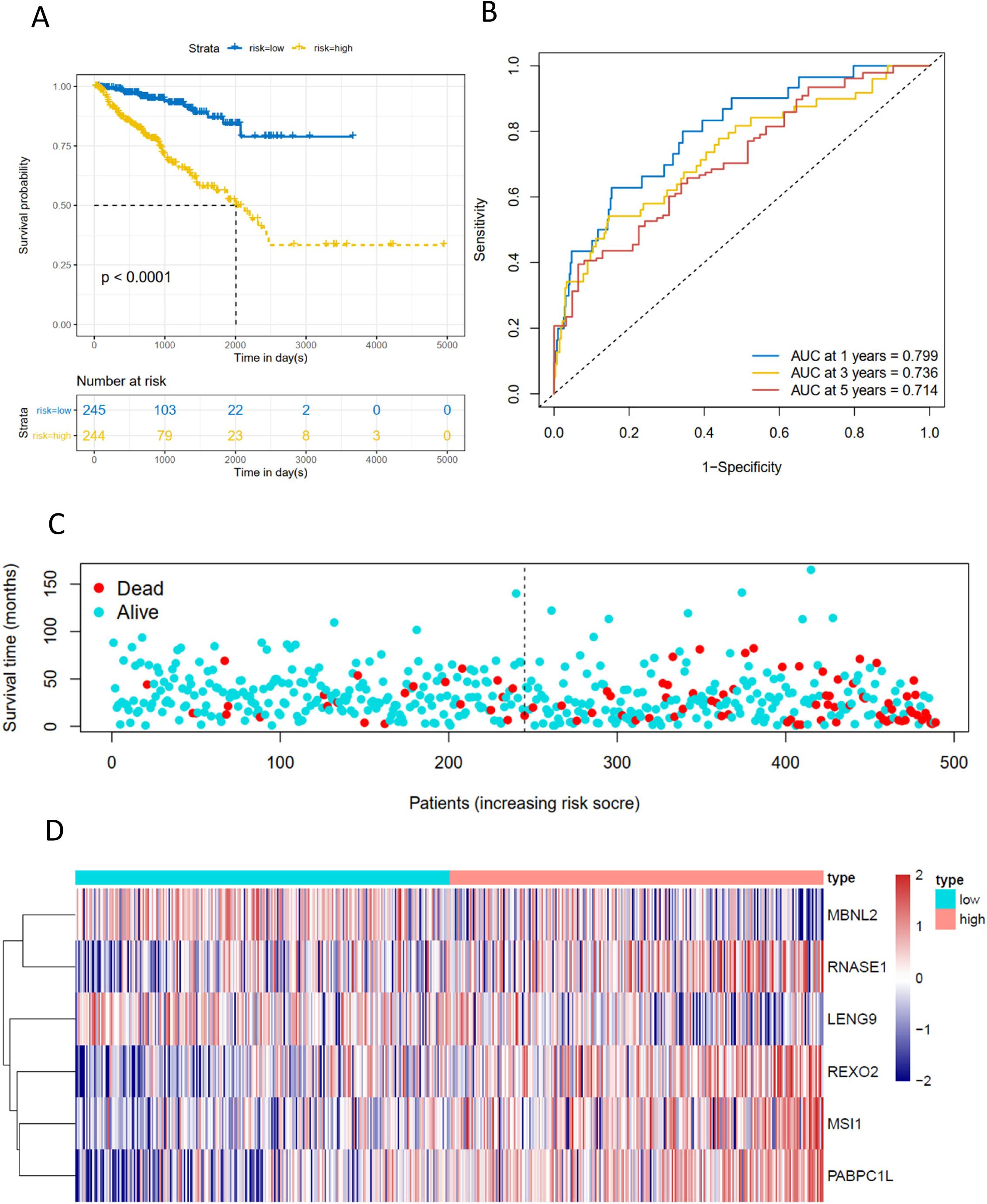
Risk score analysis of six-gene prognostic model in TCGA PRAD cohort. (A) Survival analysis according to risk score. (B) ROC analysis. (C) Survival status of patients. (D) Heatmap plot of the six genes between high and low risk group.

**Figure 6.**
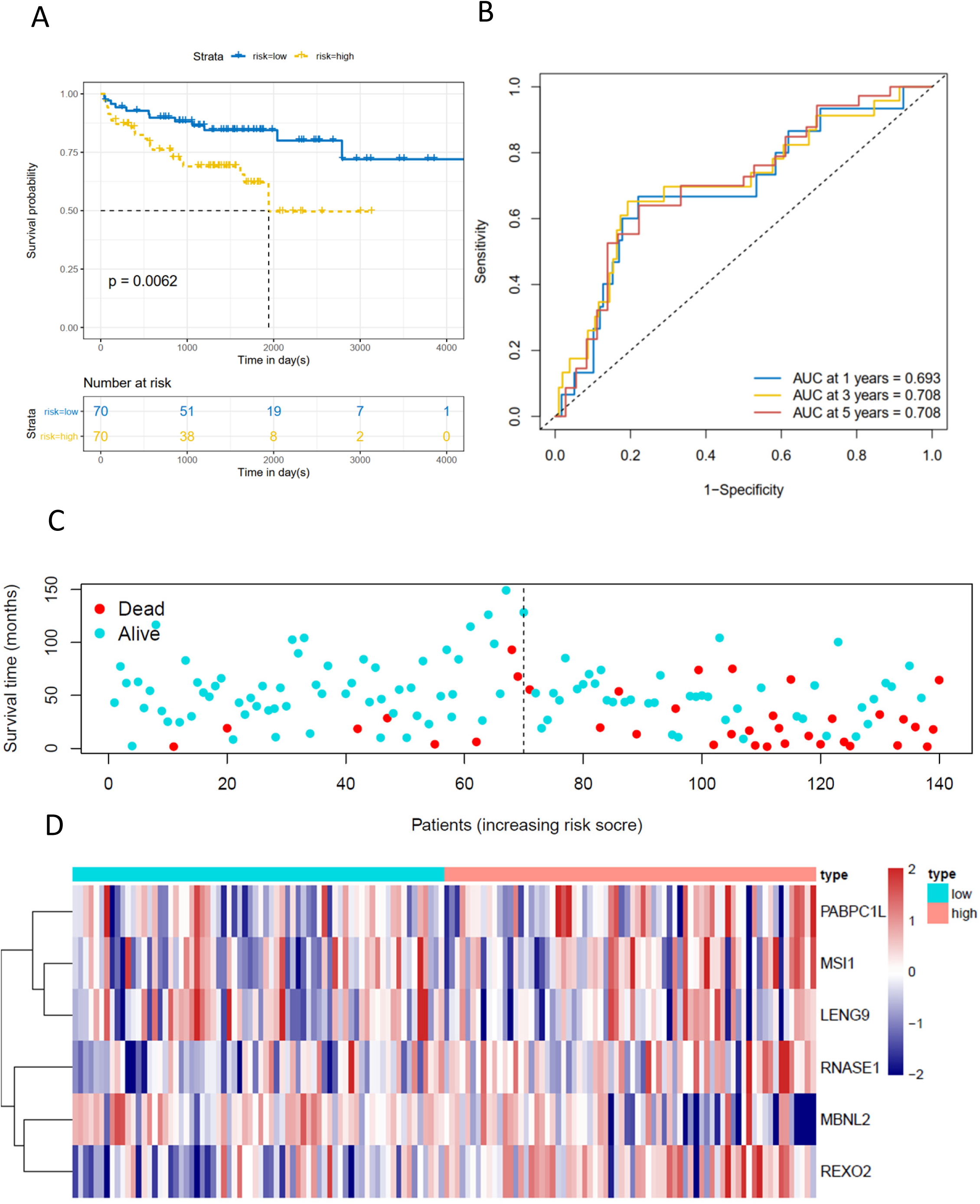
Risk score analysis of six-gene prognostic model in MSKCC cohort. (A) Survival analysis according to risk score. (B) ROC analysis. (C) Survival status of patients. (D) Heatmap plot of the six genes between high and low risk group.

### 3.3. Identificate the DEGs between high-risk and low-risk groups and explore the activated pathways

To further analysis the potential mechanism of RBPs genes, the DEGs in the different risk groups in TCGA-PRAD cohort were explored, of which 652 genes reached our threshold (adj P < 0.05, |log2FC| > 1), including 241 downregulated and 411 upregulated RBPs (**Figure 7A**). To obtain a comprehensive understanding of the relationship between risk score and the biology of PCa, we performed GO functional enrichment by using clusterProfiler package. The results showed that the DEGs were mostly associated with humoral immune response, protein activation cascade, antigen binding biology process (**Figure 7B**). Then Hallmark and KEGG functional enrichment were also performed using GSEA, which demonstrated that the high-risk group were enriched in the process of cell cycle, especially DNA repair (**Figure 7C-7D**). Therefore, we compared the mRNA expression of the key genes in the DNA damage repair (DDR) pathway among high- and low-risk group, and revealed the high expressed DDR genes in high-risk group (**Figure 7E**). Besides, the genetic alterations of many DDR key genes were also changed from low to high, such as POLE, FANCA (**Figure 7F**). These results suggested that RBPs may regulate the tumor cells biology function by changing the expression of DDR genes.

**Figure 7.**
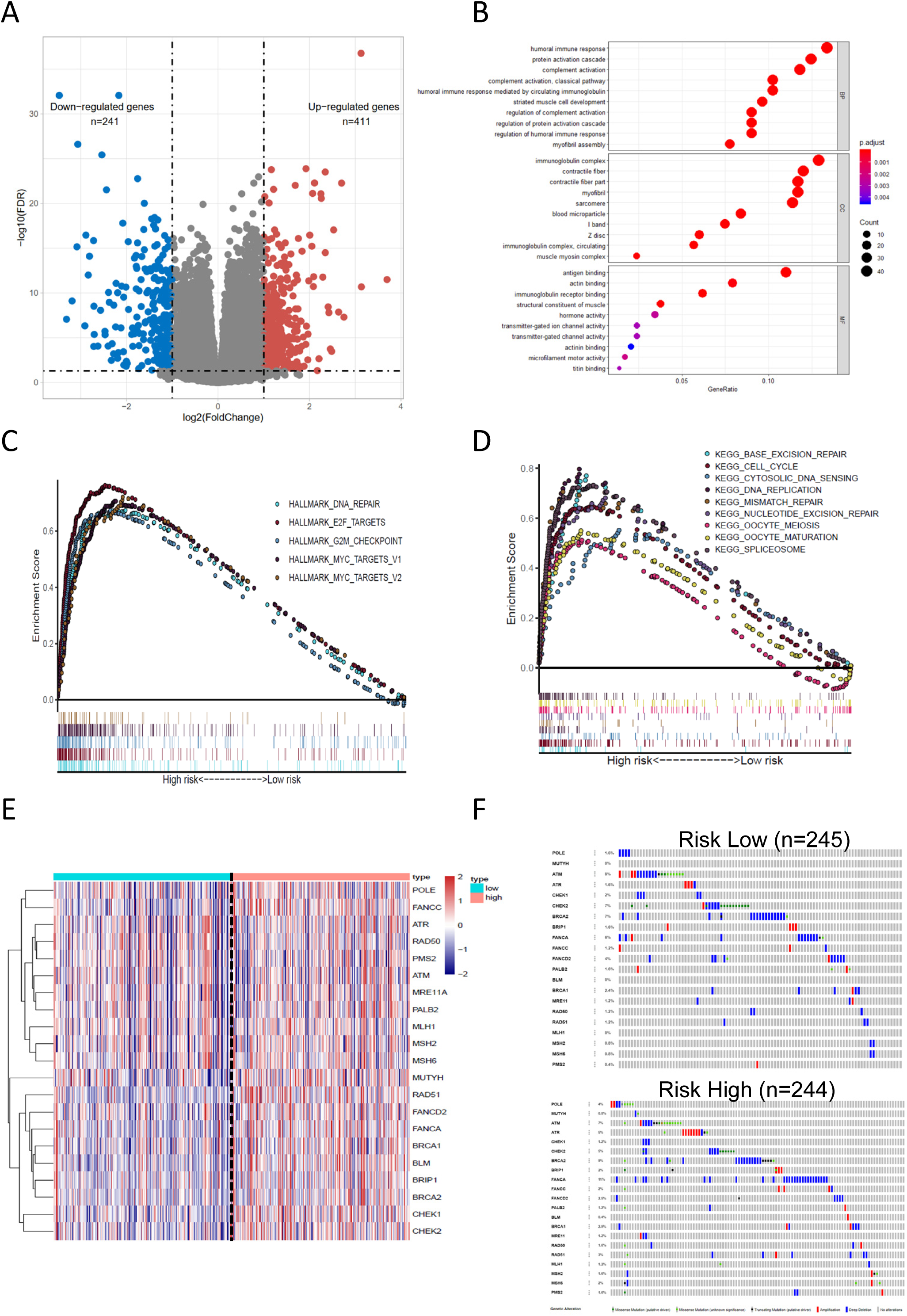
Potential biological pathways affected by RBPs. (A) The DEGs between the high-risk and low-risk groups. (B) The Gene Ontology (GO) enrichment of DEGs. (C) The Hallmark enrichment of high and low risk groups by GSEA method. (D) The KEGG enrichment of high and low risk groups by GSEA method. (E) The 21 DDR core genes mRNA expression between high and low risk group. (F) The 21 DDR core genes genetic alterations between high and low group.

### 3.4. The differences of gene copy number alteration, tumor mutant burden, cancer stemness index and sensitivity to immuno-/chemotherapy among the high- and low-risk groups

RBPs could regulate the gene transcription, especially post-transcriptional regulation, which may also have effect on the genetic alteration, such as somatic mutations, copy number alteration. Here, we further carry out analyses to get more deep understanding of the RBP risk groups. For the CNA status, the results showed that the high-risk group had the high amplification (*P*_amp_ = 0.015, *P*_amp_ = 0.00057) and deletion (*P*_del_ = 1.8e-06, *P*_del_ = 1.5e-08) in both arm and focal levels (**Figure 8A-B**). What’s more, high TMB was also found in high risk group (*P* = 1.1e-6) (**Figure 8C**). As to the tumor stemness, the stemness index in high risk group is increased, though there was no significant (**Figure 8D**). Considering TMB as a promising biomarker for immunotherapy response[34] and chemotherapy is the common method of treating advanced PCa[35]. Therefore, the immunotherapy response and different responses for cisplatin, docetaxel and bicalutamide between high risk group and low risk group were calculated based on the TIDE and GDSC database. There was no significant of immunotherapy (**Figure 8E**, *P* = 0.144), however, the estimated IC50 values showed that low risk group had a better response for Cisplatin (*P* = 3.48e-02) and bicalutamide (*P* = 7.93e-06) (**Figure 8E**).Taken together the above results, our results demonstrated that the RBPs risk model is associated with significantly higher copy number alterations, higher TMB, aberrantly drug responses, but not with cancer stemness and immunotherapy response.

**Figure 8.**
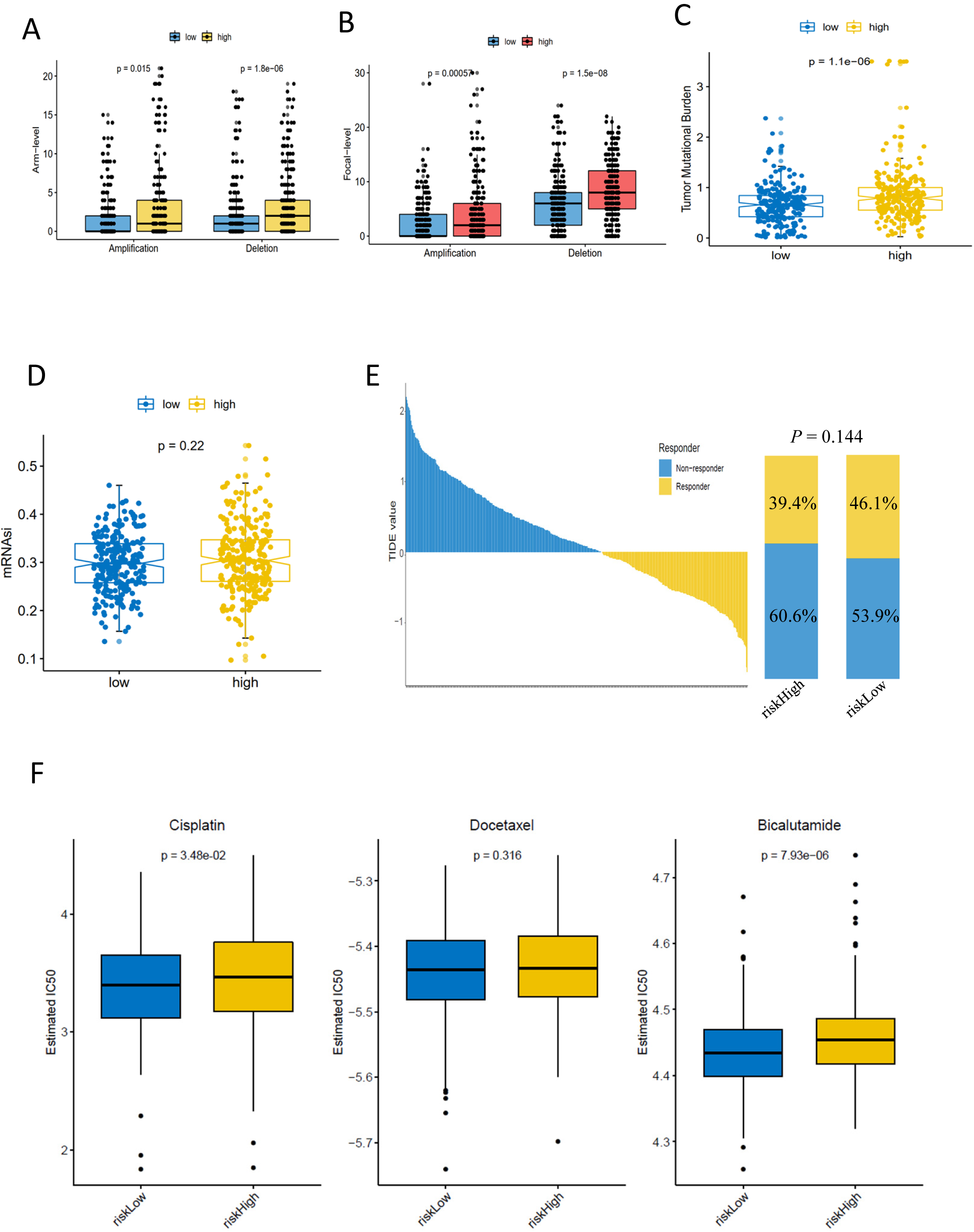
The relationship of RBP risk group with copy number alterations, tumor mutation burden, tumor stemness and immuno-/chemotherapy response. (A) Arm-level copy number amplification and deletion. (B) Focal-level copy number amplification and deletion; (C) Tumor mutant burden difference. (D) Tumor stemness difference represented by the mRNAsi. (E) Immunotherapy response based on TIDE website. (F) estimated IC50 indicates the efficiency of chemotherapy to RBP high and low risk groups by cisplatin, docetaxel and bicalutamide.

### 3.5. Related small candidate drugs screening

To identify the small molecule drugs which could treat the patients in the RBPs high risk group, CMap database was employed. Based on the 652 DEGs between high-risk group and low-risk group, the small molecule drugs with highly significant correlations were predicted (Table 2). Finally, three small molecule drugs, ribavirin, carmustine, carbenoxolone, were filter based on the enrichment (>0.6) and *P* value (< 0.05).

## 4. Discussions

RBPs aberrantly expressions had been found in several malignant tumors[20, 36],. However, little has been investigated about the expression and potential roles of RBPs in PCa. In our study, 144 differently expressed RBPs were identified in tumor compared to normal tissues based on the RNA-seq data from TCGA-PRAD cohort. Then univariate Cox regression analyses, LASSO regression analyses, multiple stepwise Cox regression analysis, and ROC analyses of hub RBPs to further explore their potential prognostic value to clinical outcome. Finally, a risk model which could predict PCa prognosis based on six RBP genes was established. These results may promote to get the new biomarkers for the diagnosis and prognosis prediction of patients with PCa.

The six hub RBPs consist of MSI1, MBNL2, LENG9, REXO2, RNASE1, PABPC1L which were associated with progression and prognosis of many cancers. MSI1, musashi1, functions as a critical protein in sensory organ precursor cells[37], which could directly binds to p21, p27, p53 mRNAs and accelerates phase transition from G0/G1 to S phase [38]. MSI1 also could control the growth of cancer by modulating the Notch, Wnt and Akt signaling pathways[39], and even be used as cancer stem cell marker to islolate adult stem cells in intestinal epithelium[40]. MBNL2, muscleblind like splicing regulator 2, which was initially found in Drosophila melanogaster[41], could regulate the expression and alternative splicing of hypoxia response of cancer cells[42]. LENG9, Leukocyte Receptor Cluster Member 9, belongs to the CCCH zine finger family, acts as an negative regulator of macrophage activation[43]. REXO2, regulate short mitochondrial RNAs generated by mtRNA processing and prevent the accumulation of double-stranded RNA[44], was reported associated with the development of inflammatory bowel disease and colorectal cancer[45]. RNASE1, which could regulate the degradation of double-stranded RNA[46], was associated with bladder cancer[47] and metastatic lesions in gastric cancer[48]. PABPC1L, binding to the polyA tails of mRNAs to regulate mRNA stability and translation[49], could reduce the cell proliferation through AKT pathway[50, 51]. Few of them has been reported in PCa. In our study, these six genes were identified aberrantly expression and associated with disease free survival. Besides, the genetic alteration of these six genes were not very high and low correlation with mRNA level. Intriguingly, the mRNA levels of MBNL2, REXO2, RNASE1 were negatively related with the methylation status (R = −0.470, R = −0.640, R = −0.486), which suggests that these three genes may be methylation-drive genes.

Next, based on the expression of these six genes, a risk model was constructed to predict the outcome of PCa in the TCGA cohort. The K-M plot showed that high risk group was dramatically associated with poorer RFS. The ROC curve analysis showed that these six genes signature with the better prognostic value to distinguish the PCa patients with poor RFS. Moreover, we also used the MSKCC database to assess the prognostic value of the six gene signature, the results were basically agreed with the results of TCGA cohort.

In order to explore the molecular mechanism of these six RBPs contributes to prostate carcinogenesis, the PCa patients were divided into two group based on the risk score. Total 652 DEGs were identified between high risk group and low risk group, then GO, KEGG and Hallmark analysis were performed, the results demonstrated that high risk group were closely related with DNA damage and repair pathway (Figure 7C-7F). Clinical and preclinical studies have revealed that DNA damage response pathways play critical role in the progression of PCa, especially in advantage stage PCa patients[52, 53]. In our study, high risk group were associated with aberrantly expression of hub gene of DDR pathway, which suggest that RBPs may regulate the DDR pathway activation and then control the biology behavior of PCa. Considering the effects exerted by RBPs and functions of DDR, the DNA copy number alteration, TMB, stemness were also analysis to explore whether RBPs are associate with genetic alteration, stemness. The results showed that the high-risk group were closely associated with amplification, deletion of arm and focal (Figure 8A-8B). The TMB was also increased in high risk group, but no significance was found in mRNAsi level (Figure 8D), which strongly reminder that RBPs may also cause the genetic alteration. Genetic alteration, especially TMB could result in the different response for drugs[54]. Then the responses for Cisplain, Docetaxel, Bicalutamide between the two groups were analysis based on the GDSC database. Our results demonstrated that high risk group had high estimated IC50, which suggested RBPs may play critical role in the response for drugs.

In addition, CMap database was used to predict some small molecule drugs with potential therapeutic efficacy for high risk group. Total 3 drugs, ribavirin, carmustine, carbenoxolone, were predicted to have anti-cancer effects in PCa. Ribavirin, known to inhibit mRNA translation, has demonstrated clinical activity of Phase I study in breast cancer, PCa, and other solid tumors[55, 56]. Carmustine attacks cancer cells through alkylation, has been used to treat lymphomas[57], brain tumors[58]. Carbenoxolone, an inhibitor of 11β-HSD, could induce apoptosis and growth inhibition in several types of cancerous cells[59-61]. These results prompted that the prognostic model of six-genes signature has a certain value in individual treatment plans of PCa patients.

Overall, our risk model built on six hub RBPs has better prognostic value for PCa patients. Moreover, the RBPs-associated gene signature was closely associated with DDR pathway, copy number alteration, TMB and drug response, suggesting that they can potentially be helpful for clinical individual therapy. Nonetheless, there are some limitations in our study. Firstly, our prognostic model was only established on the data from TCGA and MSKCC cohort, which need to be validated in clinical patient cohort and multicenter and prospective study. Secondly, further studies, including in vitro and in vivo experiments are required to clarify the molecular mechanisms for clinical practice.

In summary, we extensively explored the expression and potential prognostic abilities of differently expressed RBPs in PCa. The risk model of six RBPs was constructed, which could provide the favorable prognostic value based on these six genes expression levels. Furthermore, we found high risk group is associated with DDR pathway, copy number alteration, TMB and drug responses. Finally, three candidate small molecule drugs were predicted that had potential for RBPs high risk group, which provide directions for PCa tumors targeted therapy.

## 5. Conclusions

In our study, a prognostic-associated risk model based on six RBPs were constructed, which were closely related with DDR pathway, CNV, TMB and drug responses. These results could provide new perspective for the individual treatment of PCa patients.

## Data accessibility

All data about TCGA and MSKCC dataset are publicly available, and appear in the submitted article.

## Authors’ Contributions

XW and SG had full access to all the data in the study and take responsibility for the integrity of the data and the accuracy of the data analysis, study concept, design, and supervision. LG and JM acquisition of data and drafting of the manuscript. YZ and JG analysis and interpretation of data. ZH critical revision of the manuscript for important intellectual content. LG and JM statistical analysis. ZH, XW, SG and YZ obtaining funding, administrative, technical, or material support.

## Acknowledgments

This study was sponsored by the National Natural Science Foundation of China (No. 81972411; No. 81672555), the Young Scientists Fund of the National Natural Science Foundation of China (No. 81802544), and Young Scientists Fund of Changzhou No.2 People’s Hospital (No.2018K009).

## Conflicts of interest statement

No competing financial interests exist.

## Notes

### Competing Interest Statement

The authors have declared no competing interest.

